# Identifying tissues implicated in Anorexia Nervosa using Transcriptomic Imputation

**DOI:** 10.1101/265017

**Authors:** Laura M. Huckins, Amanda Dobbyn, Whitney McFadden, Douglas Ruderfer, Weiqing Wang, Eric Gamazon, Virpi Leppä, Roger Adan, Tetsuya Ando, Jessica Baker, Andrew Bergen, Wade Berrettini, Andreas Birgegård, Claudette Boni, Vesna Boraska Perica, Harry Brandt, Roland Burghardt, Matteo Cassina, Carolyn Cesta, Maurizio Clementi, Joni Coleman, Roger Cone, Philippe Courtet, Steven Crawford, Scott Crow, James Crowley, Unna Danner, Oliver Davis, Martina de Zwaan, George Dedoussis, Daniela Degortes, Janiece DeSocio, Danielle Dick, Dimitris Dikeos, Monika Dmitrzak-Weglarz, Elisa Docampo, Karin Egberts, Stefan Ehrlich, Geòrgia Escaramís, Tonu Esko, Xavier Estivill, Favaro Angela, Fernando Fernández-Aranda, Manfred Fichter, Chris Finan, Krista Fischer, Lenka Foretova, Monica Forzan, Christopher Franklin, Héléna Gaspar, Fragiskos Gonidakis, Philip Gorwood, Gratacos Monica, Sébastien Guillaume, Yiran Guo, Hakon Hakonarson, Katherine Halmi, Konstantinos Hatzikotoulas, Joanna Hauser, Johannes Hebebrand, Sietske Helder, Judith Hendriks, Beate Herpertz-Dahlmann, Wolfgang Herzog, Christopher Hilliard, Anke Hinney, James Hudson, Julia Huemer, Hartmut Imgart, Hidetoshi Inoko, Susana Jiménez-Murcia, Craig Johnson, Jenny Jordan, Anders Juréus, Gursharan Kalsi, Debora Kaminska, Allan Kaplan, Jaakko Kaprio, Leila Karhunen, Andreas Karwautz, Martien Kas, Walter Kaye, James Kennedy, Martin Kennedy, Anna Keski-Rahkonen, Kirsty Kiezebrink, Youl-Ri Kim, Kelly Klump, Gun Peggy Knudsen, Bobby Koeleman, Doris Koubek, Maria La Via, Mikael Landén, Robert Levitan, Dong Li, Paul Lichtenstein, Lisa Lilenfeld, Jolanta Lissowska, Pierre Magistretti, Mario Maj, Katrin Mannik, Nicholas Martin, Sara McDevitt, Peter McGuffin, Elisabeth Merl, Andres Metspalu, Ingrid Meulenbelt, Nadia Micali, James Mitchell, Karen Mitchell, Palmiero Monteleone, Alessio Maria Monteleone, Preben Mortensen, Melissa Munn-Chernoff, Benedetta Nacmias, Ida Nilsson, Claes Norring, Ioanna Ntalla, Julie O'Toole, Jacques Pantel, Hana Papezova, Richard Parker, Raquel Rabionet, Anu Raevuori, Andrzej Rajewski, Nicolas Ramoz, N. William Rayner, Ted Reichborn-Kjennerud, Valdo Ricca, Stephan Ripke, Franziska Ritschel, Marion Roberts, Alessandro Rotondo, Filip Rybakowski, Paolo Santonastaso, André Scherag, Ulrike Schmidt, Nicholas Schork, Alexandra Schosser, Jochen Seitz, Lenka Slachtova, P. Eline Slagboom, Margarita Slof-Op ’t Landt, Agnieszka Slopien, Tosha Smith, Sandro Sorbi, Eric Strengman, Michael Strober, Patrick Sullivan, Jin Szatkiewicz, Neonila Szeszenia-Dabrowska, Ioanna Tachmazidou, Elena Tenconi, Laura Thornton, Alfonso Tortorella, Federica Tozzi, Janet Treasure, Artemis Tsitsika, Konstantinos Tziouvas, Annemarie van Elburg, Eric van Furth, Tracey Wade, Gudrun Wagner, Esther Walton, Hunna Watson, D. Blake Woodside, Shuyang Yao, Zeynep Yilmaz, Eleftheria Zeggini, Stephanie Zerwas, Stephan Zipfel, Alfredsson Lars, Andreassen Ole, Harald Aschauer, Jeffrey Barrett, Vladimir Bencko, Laura Carlberg, Sven Cichon, Sarah Cohen-Woods, Christian Dina, Bo Ding, Thomas Espeseth, James Floyd, Steven Gallinger, Giovanni Gambaro, Ina Giegling, Stefan Herms, Vladimir Janout, Antonio Juliá, Lars Klareskog, Stephanie Le Hellard, Marion Leboyer, Astri J. Lundervold, Sara Marsal, Morten Mattingsdal, Marie Navratilova, Roel Ophoff, Aarno Palotie, Dalila Pinto, Samuli Ripatti, Dan Rujescu, Stephen Scherer, Laura Scott, Robert Sladek, Nicole Soranzo, Lorraine Southam, Vidar Steen, Wichmann H-Erich, Elisabeth Widen, Bernie Devlin, Solveig K. Sieberts, Nancy Cox, Hae Kyung Im, Gerome Breen, Pamela Sklar, Cynthia Bulik, Eli A. Stahl

## Abstract

Anorexia nervosa (AN) is a complex and serious eating disorder, occurring in ~1% of individuals. Despite having the highest mortality rate of any psychiatric disorder, little is known about the aetiology of AN, and few effective treatments exist.

Global efforts to collect large sample sizes of individuals with AN have been highly successful, and a recent study consequently identified the first genome-wide significant locus involved in AN. This result, coupled with other recent studies and epidemiological evidence, suggest that previous characterizations of AN as a purely psychiatric disorder are over-simplified. Rather, both neurological and metabolic pathways may also be involved.

In order to elucidate more of the system-specific aetiology of AN, we applied transcriptomic imputation methods to 3,495 cases and 10,982 controls, collected by the Eating Disorders Working Group of the Psychiatric Genomics Consortium (PGC-ED). Transcriptomic Imputation (TI) methods approaches use machine-learning methods to impute tissue-specific gene expression from large genotype data using curated eQTL reference panels. These offer an exciting opportunity to compare gene associations across neurological and metabolic tissues. Here, we applied CommonMind Consortium (CMC) and GTEx-derived gene expression prediction models for 13 brain tissues and 12 tissues with potential metabolic involvement (adipose, adrenal gland, 2 colon, 3 esophagus, liver, pancreas, small intestine, spleen, stomach).

We identified 35 significant gene-tissue associations within the large chromosome 12 region described in the recent PGC-ED GWAS. We applied forward stepwise conditional analyses and FINEMAP to associations within this locus to identify putatively causal signals. We identified four independently associated genes; *RPS26, C12orf49, SUOX*, and *RDH16.* We also identified two further genome-wide significant gene-tissue associations, both in brain tissues; *REEP5*, in the dorso-lateral pre-frontal cortex (DLPFC; p=8.52×10^−07^), and *CUL3*, in the caudate basal ganglia (p=1.8×10^−06^). These genes are significantly enriched for associations with anthropometric phenotypes in the UK BioBank, as well as multiple psychiatric, addiction, and appetite/satiety pathways. Our results support a model of AN risk influenced by both metabolic and psychiatric factors.

## Introduction

Anorexia nervosa (AN) is a serious neuropsychiatric disorder presenting with low body weight, a fear of weight gain or behaviours that interfere with weight gain, and a lack of recognition of the seriousness of the illness-. AN has the highest mortality rate of any psychiatric disorder^1^, and ranks among the leading cause of disability in young women worldwide. Despite this, little is known about the biological mechanisms underlying AN development, and few effective therapies and medications are available.

Findings from genetic and epidemiological research have encouraged broadening our conceptualization of the aetiology of AN beyond purely psychiatric causes to incorporate metabolic and other somatic factors in risk models. Recently, genome-wide association studies have revealed the first significantly associated genomic locus for anorexia nervosa^2^, as well as a number of promising sub-threshold associations^3–5^, and intriguing pathway associations. Results have implicated genes with both psychiatric and metabolic relevance, while polygenic risk score analyses and LD-Score approaches have revealed significant genetic overlap with psychiatric, metabolic and autoimmune diseases, as well as anthropometric traits.

The research findings underscore clinical observations as individuals with AN have an uncanny ability to reach and maintain extraordinarily low body mass indices (BMI) and after successful renourishment, their bodies often quickly revert to what may be an abnormally low set point^2^. Other observations include that individuals with AN tend to find eating aversive, and feelings of fullness unpleasant; dieting, restricting, and binge-purge behaviours tend to alleviate uncomfortable or painful associations with fullness in these individuals and reduce anxiety^6^. Although aversion to fullness and low appetite could be driven by dysfunction of neurobiological satiety pathways or altered levels of orexigenic hormones^7^, it is also possible that specific metabolic or gastric dysfunction enables and perpetuates dieting behaviours.

Transcriptomic Imputation (TI) provides an opportunity to test the involvement of metabolic, endocrine, adipose, and gastrointestinal (GI) tissues, as well as brain tissues, in the development of AN. These approaches leverage well curated eQTL panels to create predictors of genetically regulated gene expression (GREX)^8–10^. These predictors may be applied to large groups of genotyped individuals, to identify case-control associations with predicted differential gene expression. This approach circumvents many of the complications inherent in traditional transcriptomic analysis; for example, the need to collect large number of inaccessible tissues, which is particularly complicated in studies of early-onset psychiatric disorders^11^. Further, the prediction of genetically-regulated gene expression means that there is no ambiguity in direction of effect; unlike in RNA-seq studies, where changes in gene expression may result from medication, diet, exercise, or environmental exposures, genetically regulated gene expression necessarily precedes disease onset^8^.

An intriguing aspect of transcriptomic imputation is the opportunity to calculate predicted gene expression in a tissue-specific manner, and to use this to further inform our understanding of disease aetiology. In this study, we used gene expression predictor models for 13 brain regions (derived from CMC^12,13^ and GTEX^8,14^ data), as well as fifteen gastrointestinal, endocrine, and adipose tissues, and compared patterns of gene expression changes between cases and controls. We identified 37 significant gene-tissue associations, constituting eleven independent signals. These genes together explained 2.38% of the phenotypic variance in our study, including substantial proportions of variance explained by genes in brain tissues (51.5%), gastrointestinal tissues (16.01%), endocrine (18.6%), and adipose tissues (13.9%), supporting our theory of both psychiatric and metabolic contributions to AN risk. We identify genes with intriguing patterns of association with anthropometric traits; for example, seven of our gene-tissue associations are also significantly associated with BMI, weight, and waist circumference in the UK BioBank.

## Methods

### Samples

Genotype data were obtained from the PGC-ED collection. These data included 3,495 cases and 10,982 ancestry-matched controls^2^. Detailed diagnostic criteria used are described in the PGC-ED GWAS of these data^2^. Briefly, cases include individuals with lifetime diagnoses of either AN (including both binge-purge and restrictive subtypes) or “eating disorder not other specified (EDNOS)”, AN subtype. A small number of individuals with bulimia nervosa diagnoses were also included if they also had histories of AN. Amenorrhoea was not required for diagnosis, as it does not increase diagnostic specificity^15–17^. Exclusion criteria included schizophrenia, intellectual disability, and medical and neurological conditions which may cause weight loss.

### Transcriptomic Imputation

We imputed genetically regulated gene expression (GREX) using the CommonMind Consortium derived Dorso-lateral pre-frontal cortex (CMC DLPFC) predictor database^12^, as well as GTeX-derived predictor databases including 12 brain regions, four endocrine tissue, eight gastrointestinal/digestive tissues, and subcutaneous adipose tissue^8,14^ (Table 1). We imputed GREX in all cohorts for which we had access to raw data using PrediXcan^8^.

We tested for association between GREX and case-control status in each cohort separately, using a standard linear regression test in R. We included ten principal components as covariates to correct for population stratification. Principal components were calculated from genotype data. Raw genotype-based and summary-statistics based cohorts were meta-analysed using an odds-ratio based approach in METAL^18^.

### Establishing a threshold for genome-wide significance

We applied two significance thresholds to our data. First, we applied a threshold for each tissue, correcting for the number of genes tested within that tissue (Table 1). Second, we applied a stricter, overall threshold, correcting for all genes tested across all tissues simultaneously (234,896 tests in total, p=2.31×10^−07^).

GREX is highly correlated across tissues^14,19^, and consequently the tests across different tissues are not independent. A Bonferroni correction may therefore be overly conservative, and under-estimate the true degree of association in this study.

### Identifying independent associations

We identified a number of genomic regions with multiple associations, as well as genes with significant associations across multiple tissues. In particular, we identified a very large number of gene-tissue associations (35 significant gene-tissue associations), in the same chromosome locus identified in a recent GWAS by the PGC-ED group^20^.

We applied two methods to identified independent signals in these complex genomic regions. First, in regions with a small number of associated gene-tissue pairs (<5), we used “CoCo”, an extension to GCTA-CoJo^21^. Briefly, CoCo applies the same stepwise forward conditional analysis as in GCTA-CoJo, but allows specification of a custom linkage disequilibrium (LD) or correlation matrix instead of obtaining LD from a reference panel. Here, we calculated a GREX correlation matrix used this as the correlation matrix input to CoCo.

We used FINEMAP^22^, a shotgun stochastic search algorithm which identifies and ranks plausible causal configurations for a region, to disentangle the complex gene-tissue association patterns on chromosome 12. As for CoCo, we substituted a GREX correlation matrix in place of the standard LD-matrix input file. We constructed a 95% credible set from probable configurations specified by FINEMAP in order to identify significant gene-tissue associations within the region.

Additionally, we visually inspected patterns of correlation among the 35 gene-tissue associations in the chr12 locus using the ‘heatmap.2’ function in the ‘gplots’ R package^23^, and identified distinct clusters of GREX within this heatmap using a dendrogram cut at height 4.

### Proportion of variance explained by tissue

We calculated the proportion of phenotypic variance in our study jointly explained by all genes reaching p<1×10^−04^ in our analysis. We corrected for ten principal components and study variables using a nested model.

We divided gene-tissue associations into four categories; brain, endocrine, gastrointestinal/digestive, and subcutaneous adipose tissue. We used a series of nested models to calculate the variance explained by gene-tissue associations for each category. For example, the amount of variance explained by adipose-gene associations was calculated as the difference between the variance explained by all genes, and the variance explained by all genes except those associated in adipose tissue (eqn 1).

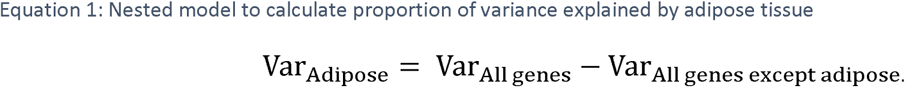

### UK BioBank analysis

We obtained publicly available GWAS summary statistics for the UK BioBank sample^24,25^. We analyzed summary statistics relating to three anthropometric traits; BMI (336,107 individuals), weight (in kg; 336,227 individuals), and waist circumference (in cm; 336,639 individuals). We obtained distributions of each trait from the UK BioBank search portal^26^ (Suppl. Table 1).

Descriptions of phenotype curation, quality control, and association models used for the UK BioBank sample are available elsewhere^25^. Briefly, quantitative traits within the sample were normalized using a rank-based inverse normal transform (INRT) prior to analysis, and analysis was carried out using a linear regression. Beta values from these associations correspond not to the ‘unit’ of the original trait (e.g., cm or kg), but to the ‘unit’ of the INRT, i.e., the standard deviation of the original trait distribution. We confirmed this by simulating distributions matching the UK Biobank traits in R, and performing an INRT on each trait.

We used MetaXcan^27^, a summary statistic based software analogous to PrediXcan, to compute gene-tissue associations for genes with p<1×10^−04^ in our prediXcan PGC-ED analysis. In order to compare association statistics between our PGC-ED and UK BioBank studies, we normalized betas to account for the variance of a gene’s GREX within each study.

### Pathway Analysis

Pathway analysis was carried out using an adaptation to MAGMA^28^. We manually assigned prediXcan genic p-values to genes in order to carry out only the gene-set enrichment analysis in MAGMA. We used Bonferroni-corrected prediXcan p-values as input for our MAGMA analyses, in three stages; first, a Bonferroni-correction for the overall best p-value for each gene across tissues; second, for the best p-value across brain regions; third, for the best p-value across non-brain tissues.

We carried out two sets of pathway analysis. First, we tested a subset of pathways for which we had prior hypotheses of involvement with psychiatric disorders^29,30^, as well as genesets related to orexigenic hormones, hunger, and satiety. Second, we carried out an agnostic pathway enrichment test including ^~^8,500 pathways obtained from publicly available databases, including KEGG^31,32^, GO^33^, REACTOME^34^, PANTHER^35,36^, BIOCARTA^37^, and MGI^38^. We included only gene sets with at least 10 genes. Gene set enrichment results from the “competitive” MAGMA analysis were used, and an FDR-correction applied within each stratum of our analysis.

## Results

### Association Tests

We calculated predicted gene expression for thirteen brain regions, four endocrine tissues, eight gastrointestinal and digestive tissue, and subcutaneous adipose tissue (derived from CMC and GTEx data^8,14,19,39^) in 3,495 cases and 10,982 controls from the PGC-ED consortium, and tested for association between predicted gene expression (GREX) and case-control status.

We identified 37 significant gene-tissue associations, and a further 22 sub-threshold associations (p<1×10^−04^; Suppl. Table 2). The majority of the significant associations (35/37) correspond to the only known genome-wide significant locus for AN^20^. We used FINEMAP^22^ to identify independent signals within this region. We identified 12 likely gene-tissue associations within this region, including four unique genes; *SUOX, RPS26, RDH16*, and *C12orf49* (Suppl. Table 3). Visual inspection (Suppl. Figure 1) and hierarchical clustering (Suppl. Figure 2) of GREX correlation patterns within this region indicate three distinct groups of associated genes, and follow our FINEMAP results closely.

We identified two additional genome-wide significant gene-tissue associations (Table 2). First, a region on chromosome two with three gene-tissue associations; increased expression of *CUL3* in the caudate basal ganglia (p=1.86×10^−06^), and increased expression of *WDFY1* and *FAM124B*, in adipose tissue (p=6.11×10^−05^, 6.73×10^−05^, respectively). We applied a stepwise forward conditional analysis in CoCo (following GCTA-COJO), using GREX correlations for all three genes (Suppl. Table 4). Neither adipose tissue association remained significant after conditioning on *CUL3-*Caudate (p=0.042, 0.25, respectively). Second, we identified decreased expression of *REEP5* in the DLPFC (p=8.34×10^−07^), and in the adrenal gland (p=6.68×10^−05^); conditioning *REEP5-* adrenal on *REEP5-*DLPFC completely ameliorates the signal (p=0.085).

Additionally, we identified 22 sub-threshold associations (p<1×10^−04^), including 17 independent associations after stepwise conditional analysis (Table 2). In particular, we identified two genes on chromosome 10 with decreased expression in the small intestine and colon (*MGMT*-small intestine, MGMT-pituitary, and *FOXI2*-colon), and two genes with increased brain expression on chromosome 17 (Supplementary table 2; *YWHAE*-hypothalamus, NTN1-nucleus accumbens).

### Comparing Tissue types

Jointly, the genetically regulated gene expression (GREX) of our 28 gene-tissue associations (p<1×10^−04^) explain 2.38% of the phenotypic variance in our study. The majority of this variance (51.5%) was explained by brain-gene associations, followed by endocrine (18.6%), gastrointestinal/digestive (16.01%), and adipose tissues (13.9%).

### Associations with anthropometry

We used publicly available GWAS summary statistics from the UK BioBank to test whether our AN associated genes were associated with anthropometric phenotypes such as BMI, weight, and waist circumference. We used a summary-statistics based approach analogous to predixcan^40^ (“MetaXcan”) to identify gene-tissue associations across all three traits, for all genes reaching p<1×10^−04^ in our analysis.

Three genes within our chromosome twelve locus were significantly associated with at least one anthropometric phenotype in the UK BioBank sample (Table 3). The direction of effect was epidemiologically consistent with our prediXcan analysis across all genes. For example, increased expression of *SUOX* in the colon, esophagus and spleen results in increased BMI (^~^0.04 BMI units/unit of gene expression; p<1.28×10^−07^), increased weight (^~^0.135kg/unit of gene expression; p<5.8×10^−08^) in the UK BioBank, and decreased risk of AN in PGC-ED (OR=0.98/unit of gene expression; p<5×10^−07^) (Figure 2A). Similarly, increased expression of *RPS26* and *RDH16* across multiple tissues is associated with increased AN risk, decreased BMI, decreased waist circumference, and decreased weight (Figure 2B).

Increased expression of *REEP5* is associated with increased weight (p<2×10^−08^) and decreased AN risk. Three sub-threshold AN genes (*BARX1, MGMT, TRIM38*) are also associated with BMI (p<x10^−13^), weight (p<2×10^−07^), and waist circumference (p=1.35×10^−08^), again with highly significant concordance of direction of effect between studies. Three sub-threshold associated genes, *BARX1, MGMT, TRIM38*, also follow this pattern of association.

This degree of shared signal and concordance of direction of effect is highly unlikely to occur by chance (binomial test p=2.39×10^−270^). Interestingly, of the seven genes within our study that are associated with BMI, weight, and waist circumference within the UK BioBank, six are associated with AN in gastrointestinal tissues. The only brain-tissue based associated gene, *REEP5*, is an olfactory gene with a potential role in taste and appetite. Although it is difficult to draw firm conclusions given the small set of genes tested and the limited sample size of our study, these results suggest that gene expression changes in metabolic tissues are more likely to have general relevance for anthropometry and weight maintenance.

### Pathway analysis

We performed pathway analyses on our AN prediXcan results across (1) all tissues, (2), brain tissues, and (3) all non-brain tissues. For each set of results, we tested 174 gene sets with prior hypotheses for involvement in psychiatric disorders, and ^~^8,500 pathways obtained from publicly available databases.

Using the best p-value across all tissues, we identified 17 significantly enriched pathways (fdr-corrected p-value<0.05; Table 4). These include multiple calcium-gated voltage channel pathways (p<0.002), axon guidance (p=1.07×10^−04^), Wnt signalling (9.93×10^−04^), the post-synaptic density (0.003), targets of the FMRP protein^41–45^ (p=0.003), as well as gene sets corresponding to neurological disease such as Alzheimer’s, Huntington’s, and Prion Disease (p<0.007). We also noted enrichment of a pathway related to circadian entrainment (p=0.0013).

Interestingly, genes involved in synthesis secretion and deacylation of ghrelin were significantly enriched within our results (p=0.0011). Examining individual genes within this pathway indicates that no single gene is driving the association; rather, the pathway includes multiple sub-threshold associations across *KLF4, BCHE, IGF1, SPCS2, ACHE, PCKS1*, and *SPSC3.* Taken together, these associations indicate lower baseline ghrelin expression in individuals with AN than in controls. For example, AN cases have lower GREX of *KLF4, SPCS2* and *SPCS3*, all of which stimulate ghrelin secretion^46^. AN cases also have increased expression of *ACHE, IGF1, PCSK1*, and *BCHE*, which inhibit ghrelin expression^47–49^. We also noted that GREX of ghrelin (GHRL) was lower in AN cases than controls across 11/12 tissues tested.

Using exclusively brain-gene association statistics as an input to our MAGMA analysis resulted in 51 significantly enriched pathways. 35/51 pathways were from the hypothesis-driven test; these included circadian entrainment (p=2.6×10^−04^), addictive behaviors (nicotine, alcohol, cocaine, and morphine dependence, p<0.0045), calcium-gated voltage channels, and a large number of pathways related to processes in the post-synaptic density (Table 4), in line with pathway results from other psychiatric disorders^10,30,50,51^. A further 25 significantly enriched pathways were identified in the agnostic analysis, including further evidence of circadian entrainment (p=1.39×10^−06^), long-term potentiation (p=4.44×10^−06^), as well as multiple pathways implicating ear and neuronal system development in mice (p<1.2×10^−04^). We noted enrichment in cyclic-AMP metabolism pathways (p<9.3×10^−05^). This pathway includes dopamine receptor gene *DRD1* (p=8.85×10^−05^), and *DRD5* (p=3.5×10^−04^), two receptors which are part of the dopaminergic pathways affected by ghrelin in the VTA and nucleus accumbens^52,53^, as well as *GCG* (Glucagon; p=1.3×10^−03^), and *APOE* (p=1.0×10^−03^) which is associated with risk for Alzheimer’s disease. CREB phosphorylation through activation of CaMKII pathway was enriched in our results (p=5.25×10^−05^). This pathway includes *AKAP9* (p=2.1×10^−04^), which regulates levels of cAMP activity in the brain, and co-localizes with NMDA receptor NR1 which in certain brain regions is involved in appetite and weight regulation^54–56^, as well as *GRIN2B* (p=5.1×10^−04^), which is associated with neurite outgrowth and risky decision making^57,58^.

Excluding brain-gene associations statistics from our pathway analysis results in only one subthreshold association (p=3.2×10^−04^; fdr-corrected p-value 0.06) in our hypothesis-driven pathway analysis, concerning circadian rhythms (albeit through a different pathway than identified in the brain-only analysis). Our agnostic pathway analysis identified only one significant association, with hyaluronic acid binding (p=2.32×10^−08^).

## Discussion

AN is a complex and serious neuropsychiatric disorder, with one of the highest mortality rates of any psychiatric disorder. As our research into the aetiology of AN develops and grows, we identify increasing levels of complexity and heterogeneity; for example, recent GWAS studies, LDScore analysis, and epidemiological evidence indicates both psychiatric and metabolic risk factors for the disorder.

Here, we used gene expression prediction models for brain, gastrointestinal/digestive, endocrine, and adipose tissues to predict genetically regulated gene expression (GREX) in 3,495 individuals with anorexia nervosa (AN) and 10,982 controls. We identified 12 independent gene-tissue associations reaching tissue-specific significance, the majority of which lie in the same chromosome 12 locus identified in a recent AN GWAS^20^. In line with our hypothesis of both psychiatric and metabolic risk having a role in AN, we identified genes with differential expression in endocrine and gastrointestinal/digestive tissues, as well as in brain.

We calculated the phenotypic variance explained by the genetically regulated expression of these 28 genes, and used a nested model to partition the variance according to tissue type. Jointly, these explain 2.38% of the phenotypic variance in our study. The majority of this variance (51.5%) was explained by brain-gene associations, followed by endocrine (18.6%), gastrointestinal/digestive (16.01%), and adipose tissues (13.9%). The proportion of variance explained by brain- and endocrine-gene associations is in line with the proportion of tests carried out in each tissue (46.3% and 16.8%, respectively). Gastrointestinal/digestive genes explain significantly less variance than we would expect given the large proportion of test performed (16.01% vs. 32.3%, binomial test, p=3.6×10^−04^), while adipose tissue-genes explain significantly more variance than we would expect (13.9% vs. 4.6%, p=2×10^−04^). This enrichment of signal within adipose tissue is of particular interest given the demonstrated overlap between adiposity and disordered eating patterns^59^, AN risk factors^60–62^, and clinical outcomes^63,64^, as well as our findings relating AN risk genes to anthropometric traits in the UK Biobank.

However, these calculations are based on the assumption that all gene-tests are independent; in fact, we note high correlation of GREX between tissues, including a large number of co-linear genes and tissues. The number of independent tests carried out is therefore likely to be substantially lower than the number of tests used in our estimate, perhaps explaining why gastrointestinal/digestive genes explain less variance than we would expect.

Among our gene-tissue associations are a number of genes which may be of particular interest. For example, decreased expression of *REEP5* in the DLPFC is associated with increased risk of AN. *REEP5* is a receptor accessory protein which promotes expression of olfactory receptors^65^. Reep5, together with RTP1 and RTP2, is required for cell surface expression of odorants, and is primarily expressed in olfactory neurons. The DLPFC has a high localized concentration of olfactory neurons, and DLPFC volume is decreased in anosmic individuals^66^. Olfaction is of particular interest in eating disorders given its role in taste and desire for food, as well as in a number of neurological disorders such as Alzheimer’s and Parkinson’s^67,68^. Individuals with AN have high rates of reported hyposmia and anosmia^67,69–72^, and perform poorly in odor discrimination tests, compared to healthy controls. Importantly, odor discrimination ability and hyposmic status correlates more strongly with BMI than with any specific disordered eating behavior, even among individuals with AN^73^. Previous studies have also demonstrated differential expression of olfactory genes following eight restoration in individuals with Anorexia Nervosa^74^. In line with this, we identified a direct correlation between *REEP5* expression and body weight in the UK BioBank; each additional unit of gene expression corresponds to ^~^140 g additional body weight, and an AN OR of 0.85. Taken together these results suggest that *REEP5* may have a general role in body size and BMI through altered olfactory cues, and may be of interest to researchers studying appetite and satiety, as well as obesity, normal variation in BMI, and AN. *REEP5* has also been implicated in major depressive disorder and antidepressant response in previous studies^75^.

We identified four significantly associated genes within our complex chromosome 12 locus. Three of these genes (*SUOX, RPS26, RDH16*) are significantly associated with AN across a range of gastrointestinal tissues (Figure 1), and have highly correlated expression across almost all non-brain tissues tested. All three of these genes are significantly correlated with anthropometric traits in the UK BioBank analysis (Figure 2), and all have consistent directions of effects with our AN prediXcan analysis: that is, the change in expression which increases body size also decreases AN risk.

**Figure 1:**
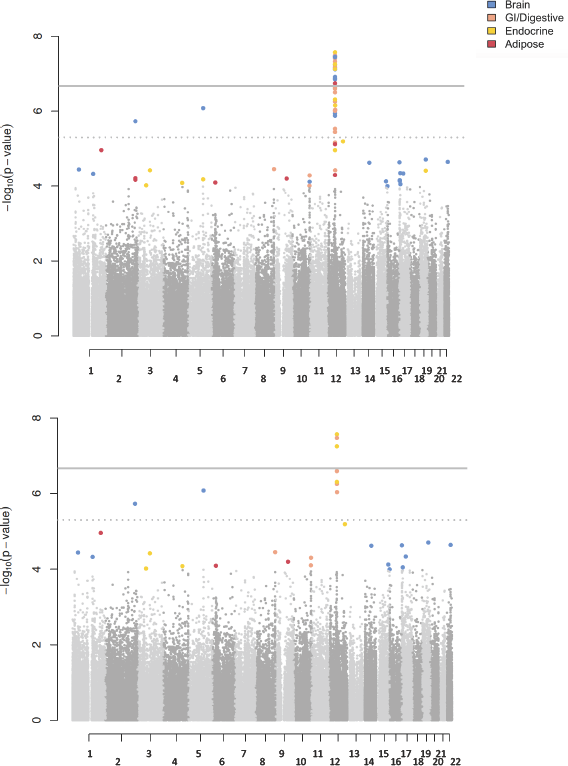
Genic associations in Anorexia Nervosa A) We identify 37 significant gene-tissue associations across brain, GI/digestive, endocrine, and adipose tissues B) 14 gene-tissue associations remain significant after applying CoCo and FINEMAP.

**Figure 2A:**
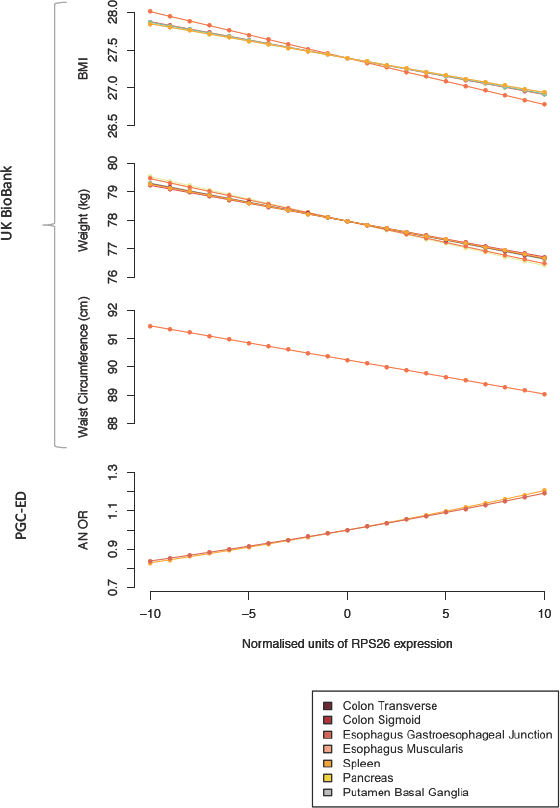
Genetically regulated expression of *RPS26* is significantly associated with BMI, weight and waist circumference in the UK BioBank, and with AN in PGC-ED

**Figure 2B:**
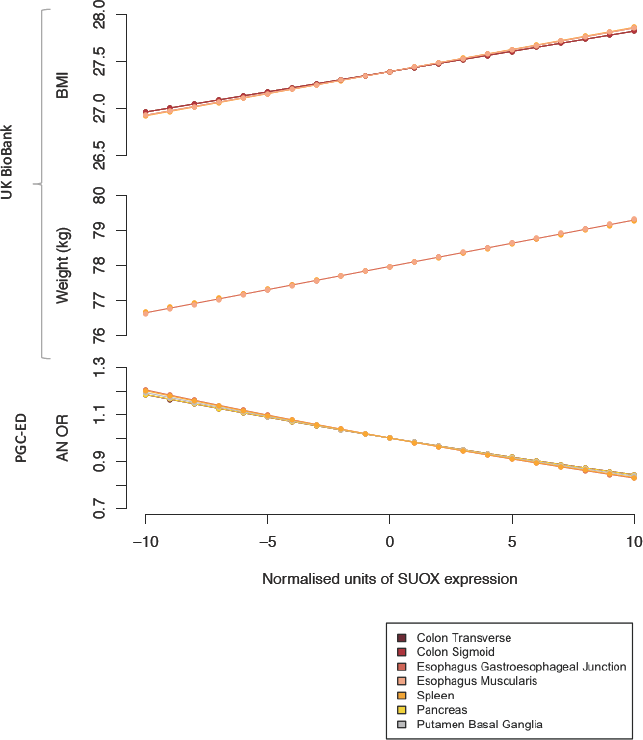
Genetically regulated expression of *SUOX* is significantly associated with BMI and weight in the UK BioBank, and with AN in PGC-ED

Little is known about the function of *C12orf49*, the fourth gene in this locus, although SNPs within the gene have also previously been associated with BMI, waist circumference, and waist-hip ratio^76^. Taken together, this evidence implies that the locus on chromosome 12 is likely to be generally associated with BMI and body size, rather than any specific eating disordered behaviours. The fine-mapping and characterization of this locus supports our hypothesis of a role for metabolic dysregulation in AN.

Increased expression of *CUL3* (Cullin 3) in the caudate basal ganglia was associated with increased risk of AN in our study (OR=1.07). Dysregulation of *CUL3* is associated with pseudohypoaldosteronism^77^, a disorder characterized by sodium imbalance in the body and often presenting with low body weight. Mutations in *CUL3* are associated with schizophrenia^78^, autism^79^ and non-response to anti-depressants^80^. Variants lying near to *CUL3* were identified in the first GWAS of AN, although these did not reach genome-wide significance^81^.

Among our subthreshold gene-tissue associations, we identified a number of genes previously associated with psychiatric^13,78^ and neurological disorders (for example, *FURIN^13, 78, 82^, ADAMTS9^83–86^, MGMT^86, 87^, SMDT1*^78^ *TMEM108^88^*), as well as with abnormal behavioural responses in knock-out mice models^38,89–91^ (*ADAMTS9, CITED4, FOXI2, FURIN, SMDT1, Ell TMEM108).* We also noted a number of genes with prior associations with anthropometric traits, both in humans (*ADAMTS9^92, 92–96^, MGMT^94, 97, 98^*) and in mice^38,89–91^ (*CITED4, FOXI2, FURIN*, *RDH16*, *SMDT1, TMEM108*), as well as genes associated with gastric and esophageal complaints (*BARX1*^99^) in humans, and abnormal defecation patterns in mice^38,89–91,100^ (*RDH16, CITED4*), and with disorders and traits known to be comorbid with AN (*TMEM108^101–104^).*

Our pathway analysis identified a large number of significantly enriched pathways. In particular, multiple pathways indicate a role for the post-synaptic density (including PSD95, targets of the FMRP protein, glutamate receptor genes, among others), which has previously been implicated in other psychiatric disorders. Four pathways are associated with addiction and addictive behaviours, including nicotine addiction, alcoholism, cocaine addiction, and amphetamine addiction. Illicit drug use is significant enriched among individuals with eating disorders, in particular AN^105^, although this is, to our knowledge, the first study identifying shared genetic risk factors.

Circadian entrainment and clock genes are highly enriched among our data. Longstanding hypotheses implicated disrupted circadian rhythms in a range of mood disorders, particularly depression and bipolar disorder^106–108^. Further, behavioural patterns in individuals with AN (for example excessive exercise^109–111^ and lack of sleep) have long provided epidemiological evidence for circadian rhythm disruption in AN. Circadian rhythms may also have a role is regulating appetite and satiety pathways^7,112,113^.

Our analysis also implicates pathways concerning taste and olfactory transduction, as well as ghrelin secretion. Ghrelin is an orexigenic hormone with a documented role in appetite and satiety^114–118^ as well as in gut motility^117–119^. Our results suggest that individuals with AN may have decreased circulating ghrelin levels due to increased genetically regulated expression of ghrelin inhibitors, and decreased GREX of Ghrelin stimulators. Ghrelin enhances appetite and increases food intake in humans; lowered baseline circulating ghrelin levels may begin to explain decreased hunger and desire for food in individuals with AN. Previous studies have documented dysregulation of ghrelin, leptin and glucagon in individuals with AN^120^. However, these studies are by definition performed after long periods of starvation or food restriction, meaning that causation is difficult to disentangle from consequences of eating disordered behaviours; it is likely that the increased ghrelin levels seen in these studies is a consequence of long-term fasting, rather than causative. In this study, we assess only genetically regulated gene expression (GREX), meaning that any associations identified are not affected by diet or environment. Instead, these results may indicate an altered “baseline” level of circulating ghrelin in individuals with AN.

There are a number of limitations that should be taken into account. First, the sample size of our study is small, especially compared to GWAS sample sizes in other psychiatric disorders^121,122^. It is likely that increasing sample size substantially will yield many new insights into the aetiology of anorexia nervosa, and that current sub-threshold associations may lose significance as sample size increases. Similarly, transcriptomic imputation approaches rely on large, well-curated reference panels in order to build GREX predictor models; here, we have used reference panels constructed from GTeX^8,14^ and CommonMind Consortium data^10,13^, including the largest collections of publicly available post-mortem brain tissues. We have shown previously that there is a significant correlation between the sample sizes used to construct these predictors and the number of genes included in each predictor database, and that a number of these databases are therefore likely underpowered^10^.

Our analysis highlights the need for greater investigation into the complex aetiology of anorexia nervosa. Transcriptomic Imputation allows us to identify significant gene-tissue associations with anorexia nervosa, and indicates an excess of signal in adipose tissue. It is clear from these results that both psychiatric and metabolic risk factors play a role in AN risk; these factors should be carefully considered in the design of future studies, as well as in how AN is perceived and considered by clinicians treating individuals with AN.

**This manuscript is dedicated to the memory of Professor Pamela Sklar.**

## Acknowledgements

Data were generated as part of the CommonMind Consortium supported by funding from Takeda Pharmaceuticals Company Limited, F. Hoffman-La Roche Ltd and NIH grants R01MH085542, R01MH093725, P50MH066392, P50MH080405, R01MH097276, RO1-MH-075916, P50M096891, P50MH084053S1, R37MH057881 and R37MH057881S1, HHSN271201300031C, AG02219, AG05138 and MH06692.

---

Brain tissue for the study was obtained from the following brain bank collections: the Mount Sinai NIH Brain and Tissue Repository, the University of Pennsylvania Alzheimer’s Disease Core Center, the University of Pittsburgh NeuroBioBank and Brain and Tissue Repositories and the NIMH Human Brain Collection Core. CMC Leadership: Pamela Sklar, Joseph Buxbaum (Icahn School of Medicine at Mount Sinai), Bernie Devlin, David Lewis (University of Pittsburgh), Raquel Gur, Chang-Gyu Hahn (University of Pennsylvania), Keisuke Hirai, Hiroyoshi Toyoshiba (Takeda Pharmaceuticals Company Limited), Enrico Domenici, Laurent Essioux (F. Hoffman-La Roche Ltd), Lara Mangravite, Mette Peters (Sage Bionetworks), Thomas Lehner, Barbara Lipska (NIMH).

The Genotype-Tissue Expression (GTEx) Project was supported by the Common Fund of the Office of the Director of the National Institutes of Health, and by NCI, NHGRI, NHLBI, NIDA, NIMH, and NINDS. The data used for the analyses described in this manuscript were obtained from the GTEx Portal on 09/05/16. BrainSpan: Atlas of the Developing Human Brain [Internet]. Funded by ARRA Awards 1RC2MH089921-01, 1RC2MH090047-01, and 1RC2MH089929-01.

UK BioBank analyses were carried out using results obtained from the Neale and Rivas Labs. Results were downloaded from the Global Biobank Engine (Global Biobank Engine, Stanford, CA (URL: http://gbe.stanford.edu) [September 30^th^, 2017]. The authors would like to thank the Rivas lab for making the resource available.

**Acknowledgements for the Eating Disorders Working Group of the Psychiatric Genomics Consortium (PGC-ED)** The Wellcome Trust Wellcome Trust WT088827/Z/09; WT098051;

Swedish Research Council (VR Dnr: 538-2013-8864);

We thank the Price Foundation for their support of recruiting patients, collecting clinical information and providing DNA samples used in this study. We also thank the Klarman Family Foundation for supporting the study. We thank the technical staff at the Center for Applied Genomics (CAG) at CHOP for generating genotypes used for analyses and the nursing, medical assistant and medical staff for their invaluable assistance with sample collection. Yiran Guo was funded by the 2011-2014 Davis Foundation Postdoctoral Fellowship Program in Eating Disorders Research Award. Dong Li was funded by the 2012-2015 Davis Foundation Postdoctoral Fellowship Program in Eating Disorders Research Award. Genome-wide genotyping for CHOP samples was funded by an Institutional Development Fund to CAG from CHOP. The study was additionally funded through the Electronic Medical Records and Genomics (eMERGE) Network (U01 HG006830) by National Fluman Genome Research Institute of National Institutes of Health, and also funded by donation from the Kurbert Family;

National Institutes of Health: K01MH093750; K01MH106675; K01MH109782; K02AA018755-06, R01AA015416-08, 5U01MH094432-04, 3U01MH094432-03S1; R01 MH109528; D0886501; R01 MH092793;

This work received grants from EC Framework V Factors in Healthy Eating, from INRA/INSERM (4M406D), and from PHRC ENDANO (2008-A01636-49);

European Commission (2008-2011) as an Early Stage Researcher from the Research Training Network INTACT (Individually Tailored Stepped Care for Women with Eating Disorders) in the Marie Curie Program (MRTN-CT-2006-035988);

Gerome Breen acknowledges support from the National Institute for Health Research (NIHR) Biomedical Research Centre at South London and Maudsley NHS Foundation Trust and King’s College London. The views expressed are those of the authors and not necessarily those of the NHS, the NIHR, or the Department of Health.

Veneto Region Grant BIOVEDA; Contract grant number: DGR 3984/08;

Academy of Finland (28327, 286028); (Dr. Raevuori, grant number 259764);

The German Ministry for Education and Research (National Genome Research Net-Plus 01GS0820 and 01KU0903), the German Research Foundation (DFG; HI865/2-1), the European Community’s Seventh Framework Programme (FP7/2007-2013) under grant agreement no. 245009 and no.262055." We thank the German Ministry for Education and Research for funding the ANTOP-study (project number 01GV0624) AS was supported by the Federal Ministry of Education and Research (BMBF), Germany, FKZ 01EO1502; German Federal Ministry for Education and Research (BMBF) 01GV0601 and 01GV0624; German Ministry for Education and Research (National Genome Research Net-Plus 01GS0820) and the German Research Foundation (DFG; HI865/2-1); MHT received grant support from the Alexander von Humboldt Foundation, the Helmholtz Alliance ICEMED-Imaging and Curing Environmental Metabolic Diseases, through the Initiative and Networking Fund of the Helmholtz Association, the Helmholtz cross-program topic “Metabolic Dysfunction,” and the Deutsche Forschungsgemeinschaft (DFG-TS226/1-and TS226/3-1) and the European Research Council Consolidator Grant (HepatpMetaboPath)” Ministry for Research and Education, Germany; The Helmholtz Alliance ICEMED-Imaging and Curing Environmental Metabolic Diseases, through the Initiative and Networking Fund of the Helmholtz Association, the Helmholtz cross-program topic “Metabolic Dysfunction,” and the Deutsche Forschungsgemeinschaft (DFG-TS226/1-1 and TS226/3-1); This work was supported by the Deutsche Forschungsgemeinschaft (EH 367/5-1 and SFB 940) and the Swiss Anorexia Nervosa Foundation;

Andrew Bergen is supported by a Professional Services Agreement with the Regents of the University of California; Resnick Family Chair in Eating Disorders; Klarman Family Foundation;

Research Council of Norway (RCN), and South-East Norway Regional Health Authority (SEN); Bergen Research Foundation, NFR (NORMENT-SFF), NCNG: This sample collection was supported by grants from the Bergen Research Foundation and the University of Bergen, the Dr Einar Martens Fund, the K.G. Jebsen Foundation, the Research Council of Norway, to SLH, VMS and TE; Supported by the Research Council of Norway (#248778, # 223273); The twin program of research at the Norwegian Institute of Public Health was supported by grants from The Norwegian Research Council and The Norwegian Foundation for Health and Rehabilitation;

China Scholarship Council (Shuyang Yao);

Genome Canada, the government of Ontario, the Canadian Institutes of Health Research, University of Toronto McLaughlin Centre; Ontario Mental Health Foundation for funding the recruitment and collection of the DNA samples. Ministry of Health of Ontario AFP Innovation Fund;

Grants 324715 and 480420 from the National Health and Medical Research Council (NHMRC) to TDW supported this work. Administrative support for data collection was received from the Australian Twin Registry, which is supported by an Enabling Grant (ID 310667) from the NHMRC administered by the University of Melbourne; Matthew Flinders Fellowship, Flinders University, South Australia, Australia

Internal Grant Agency of the Ministry of Health of the Czech Republic IGA MZ _R NT 14094- 3/2013;

Research of Korea Centers for Disease Control and Prevention Fund (code# HD16A1351);

Nicole Soranzo’s research is supported by the Wellcome Trust (Grant Codes WT098051 and WT091310), the EU FP7 (EPIGENESYS Grant Code 257082 and BLUEPRINT Grant Code†HEALTH-F5-2011-282510) and the National Institute for Health Research Blood and Transplant Research Unit (NIHR BTRU) in Donor Health and Genomics at the University of Cambridge in partnership with NHS Blood and Transplant (NHSBT). The views expressed are those of the author(s) and not necessarily those of the NHS, the NIHR, the Department of Health or NHSBT.

Psychiatry Research Trust (registered charity no. 284286);

Spanish Ministry of Economy and Competitiveness (MINECO) no. SAF2013-49108-R, the Generalitat de Catalunya AGAUR 2014 SGR-1138, the European Commission 7th Framework Program (FP7/2007-2013) 262055 (ESGI); Instituto de Salud Carlos III (FIS PI14/290 and CIBERobn

Supported by MH CZ - DRO (MMCI, 00209805);

An unrestricted grant from the Lundbeck Foundation, iPSYCH (Initiative for Integrative Psychiatric Research); and by Aarhus University for CIRRAU (Centre of Integrated Register-Based Research);

This research was supported by a ZonMW VIDI Grant (91786327) from The Netherlands Organization for Scientific Research (NWO) to Prof. dr. Martien Kas;

This study was supported by EU H2020 grants 692145, 676550, 654248, Estonian Research Council Grant IUT20-60, NIASC, EIT ñ Health and NIH-BMI Grant No: 2R01DK075787-06A1 and EU through the European Regional Development Fund (Project No. 2014-2020.4.01.15-0012 GENTRANSMED;

Ulrike Schmidt receives salary report from the National Institute of Health Research Mental Health Biomedical Research Centre at South London and Maudsley National Health Service Foundation Trust and King’s College London.

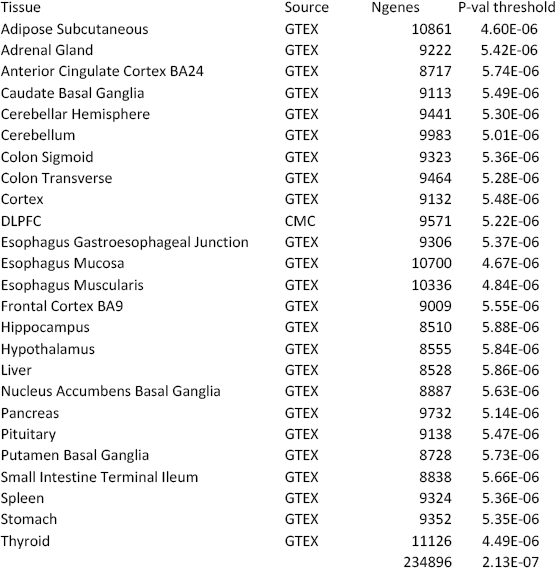

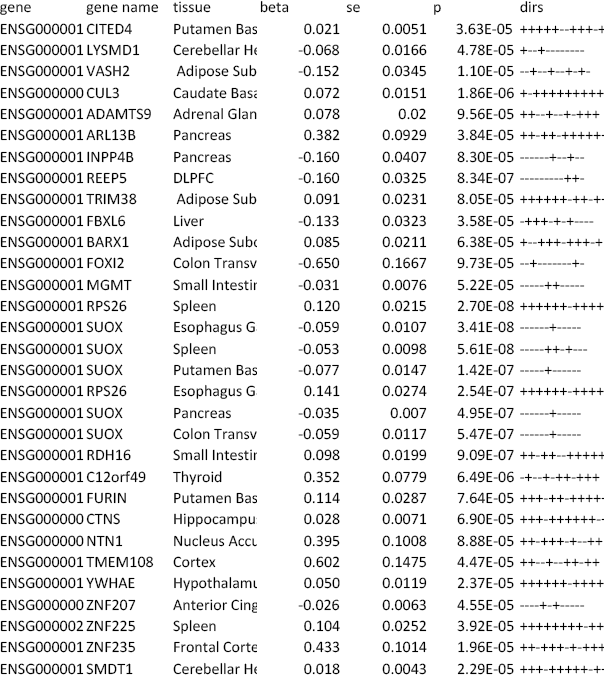

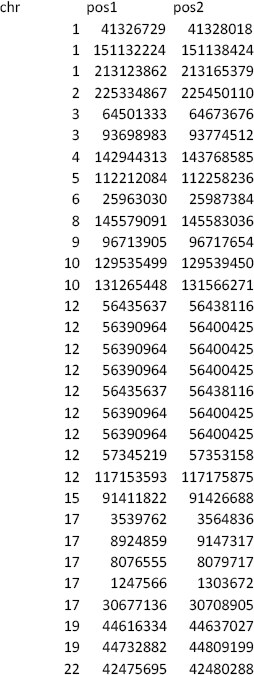

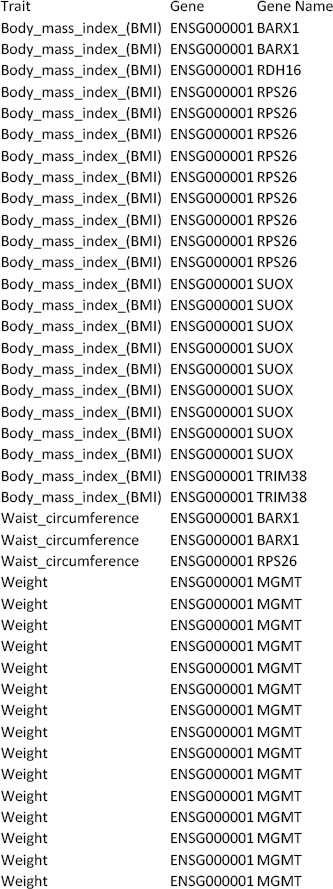

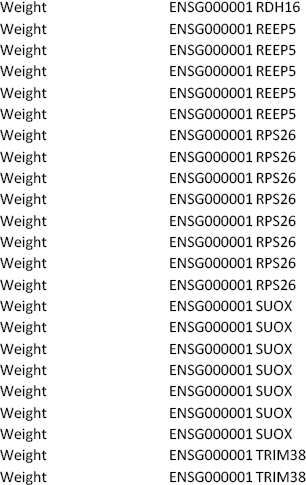

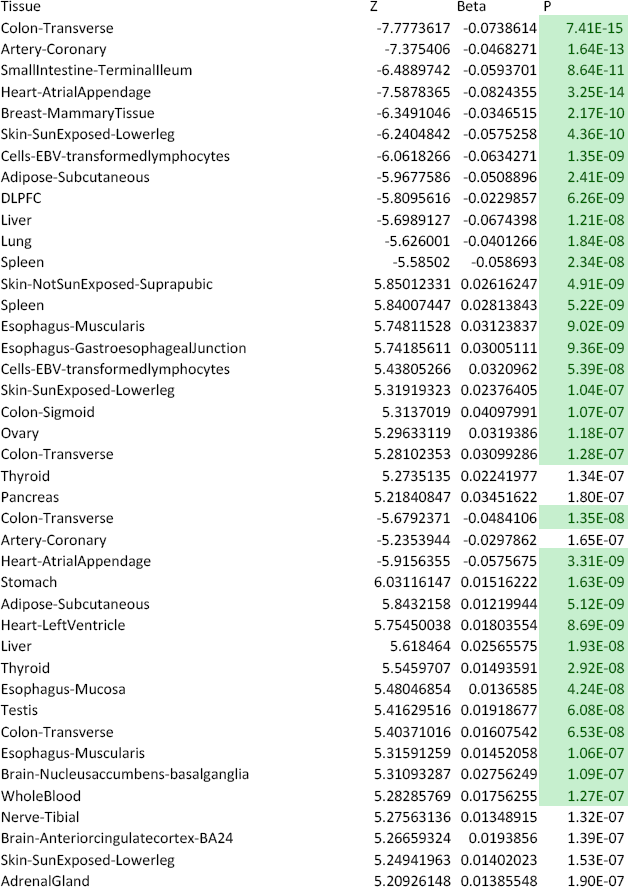

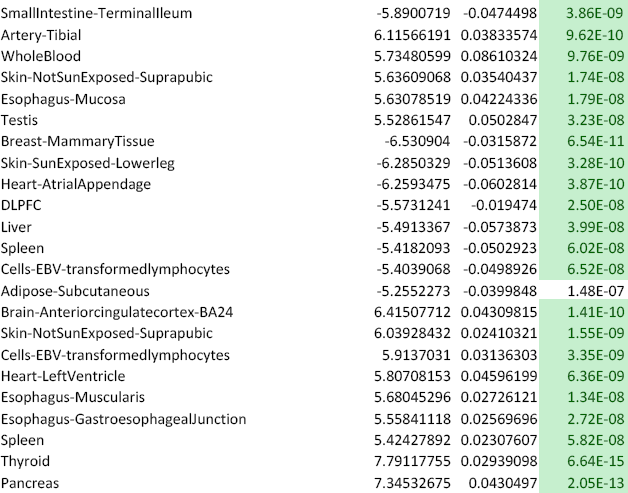

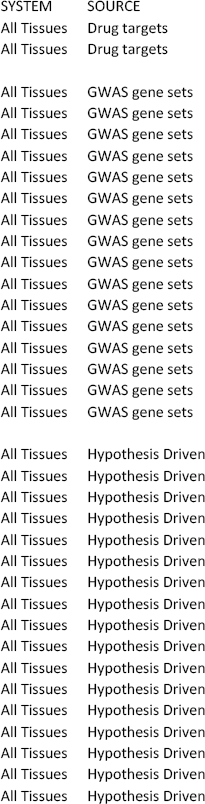

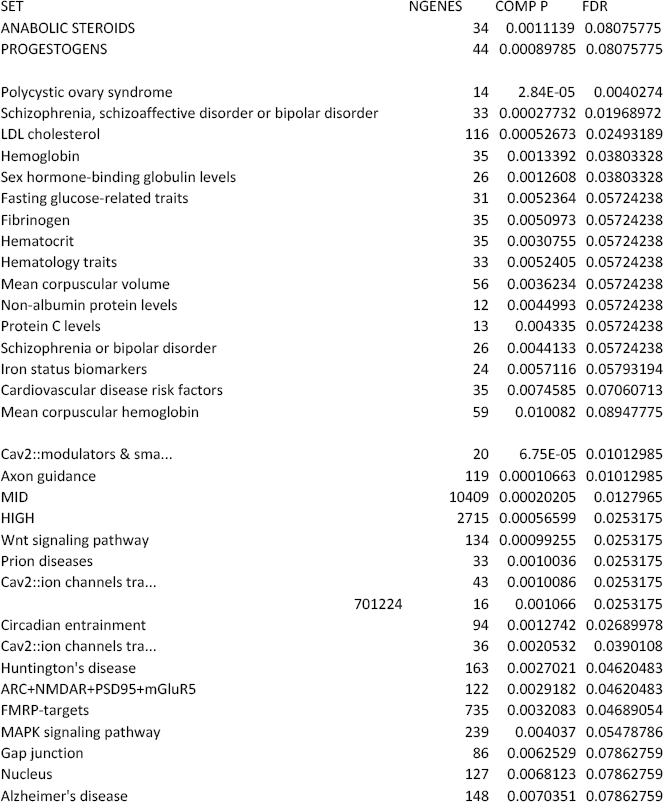

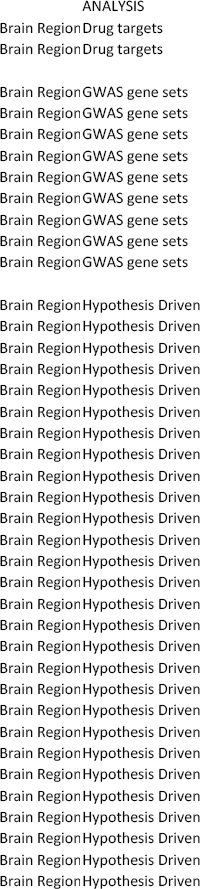

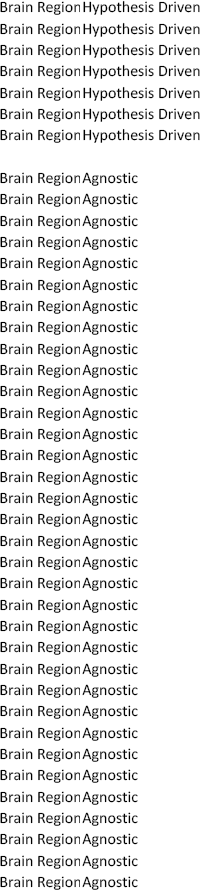

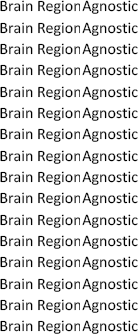

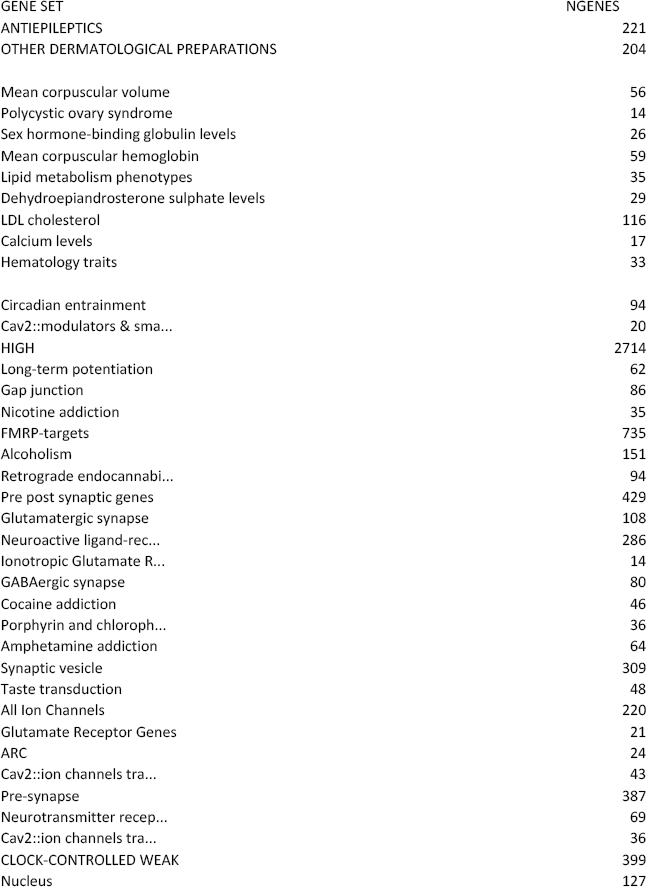

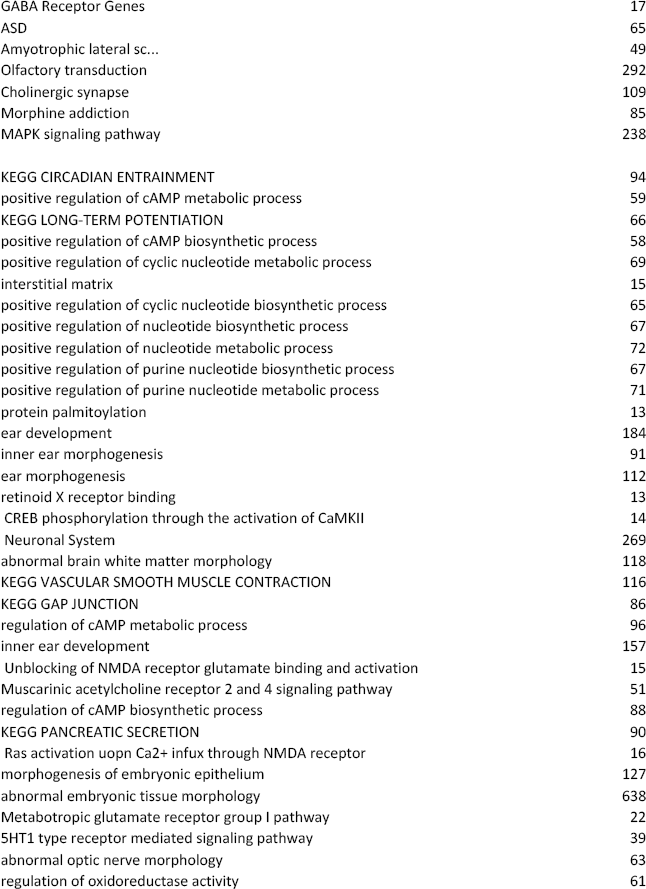

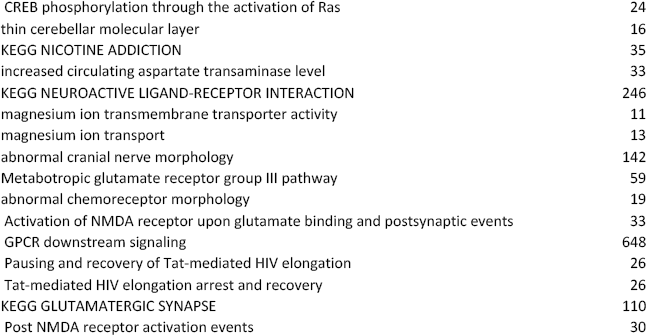

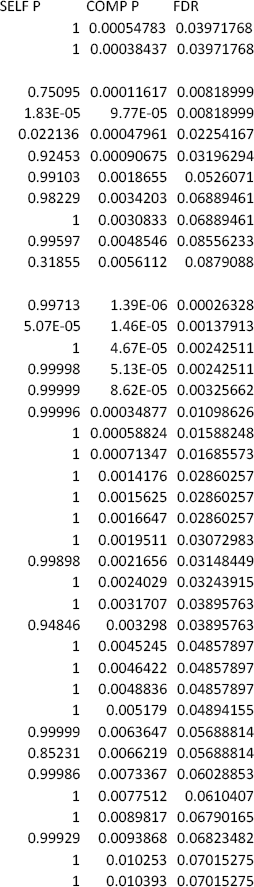

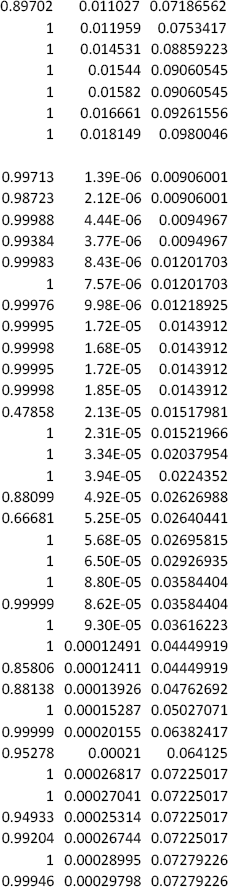

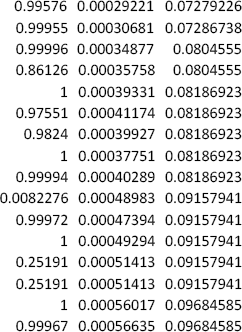

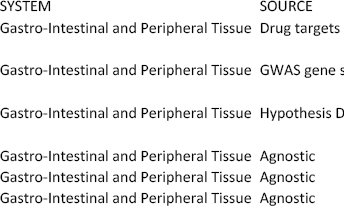

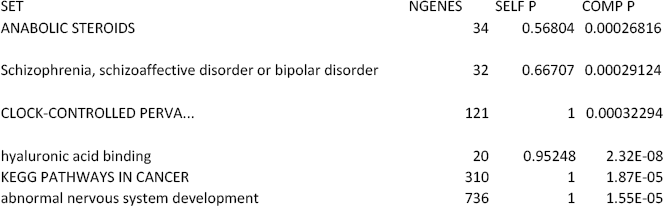

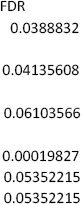

